# mmu-miR-1291 Alleviates Oxidative Damage in Myocytes by Modulating the Process of Gene-Mediated Ubiquitination

**DOI:** 10.1101/2025.04.25.649032

**Authors:** Jiameng Shen, Dongdong Bo, Yuanyuan Wang, Yueyu Bai

**Affiliations:** National Key Laboratory of Cotton Bio-breeding and Integrated Utilization, School of Agricultural Sciences, Zhengzhou University; Key Laboratory of Innovative Utilization of Local Cattle and Sheep Germplasm Resources (Co-construction by Ministry and Province), Ministry of Agriculture and Rural Affairs, Zhengzhou 450001, China

**Keywords:** Skeletal muscle, C2C12, β-sitosterol, Sodium palmitate, Oxidative damage, Myocytes, mmu-miR-1291, *Wsb1*

## Abstract

BS exhibited a significant restorative effect on PA-induced oxidative damage in C2C12 cells. And a set of miRNAs that were significantly differentially expressed under oxidative stress conditions was identified. Through target gene prediction, a key miRNA potentially involved in oxidative stress regulation was selected. Additionally, differential expression and enrichment analyses led to the identification of a series of mRNAs affected by oxidative stress. Based on miRNA target prediction, the Wsb1 gene was selected for further investigation. A dual-luciferase reporter assay confirmed that this miRNA directly regulates Wsb1. This suggests that the identified miRNA may influence the oxidative stress process by regulating Wsb1 expression and potentially be involved in modulating the NF-_**κ**_B signaling pathway.

## 1. Introduction

β-Sitosterol (BS), also known as 3β-stigmast-5-en-3-ol, 22,23-dihydrostigmasterol, rhamnol, and stigmasterol, belongs to the tetracyclic triterpenoid compounds and possesses a distinct odor. BS shares similar chemical constituents with cholesterol, featuring an ethyl group at the 24th carbon position, which distinguishes it from animal cholesterol [1,2].

β-Sitosterol, one of the most abundant dietary plant sterols, exhibits properties that alleviate various types of cancers, including lung cancer, gastric cancer, ovarian cancer, breast cancer, prostate cancer, colon cancer, and leukemia. Additionally, β-sitosterol interferes with multiple cellular signaling pathways, involving cell cycle, apoptosis, proliferation, survival, invasion, angiogenesis, metastasis, and inflammation [2-5]. It possesses a wide range of pharmacological activities, such as anti-inflammatory, antioxidant, cholesterol-lowering, hypoglycemic, and immunomodulatory effects [6,7].

Skeletal muscle, one of the most crucial locomotor organs in mammals, is composed of multinucleated muscle fibers exhibiting the typical structural features of striated muscle. Its fundamental unit is the sarcomere, which contracts through the sliding of thick and thin filaments in response to neuronal signals [8]. Skeletal muscles are generally attached to bones via tendons and are surrounded by connective tissue layers such as the epimysium, perimysium, and endomysium, which provide support, protection, and nutritional supply [9]. The interior of skeletal muscle is rich in blood vessels and neural networks, satisfying its high metabolic demands [10,11]. Besides controlling movement, skeletal muscle plays a pivotal role in posture maintenance, thermogenesis, and energy metabolism [12].

Oxidative stress arises from the excessive accumulation of reactive oxygen species (ROS) due to an imbalance between ROS generation and antioxidant capacity, which normally does not cause damage to cellular molecules such as lipid peroxidation or protein carbonylation [13]. ROS refer to highly reactive species containing oxygen-derived free radicals, which are produced by oxidase enzymes and scavenged by clearance systems involving enzymatic or non-enzymatic reactions. An imbalance between ROS production and clearance systems can lead to elevated ROS levels [14]. Skeletal muscles are rich in mitochondria, and mitochondrial ROS production is essential for maintaining muscle-related processes, including skeletal muscle development, muscle mass maintenance, mitochondrial biogenesis, and muscle repair [15]. However, excessive ROS production can damage biomolecules and cellular structures, reduce mitochondrial membrane potential, and ultimately induce apoptosis, leading to a range of diseases [16].

## 2. Materials and Methods

### 2.1. Cell Culture

C2C12 cells were purchased from Zhejiang Meisen Cell Technology Co., Ltd. (Hangzhou, China). The C2C12 cells were cultured in Dulbecco’s Modified Eagle’s Medium (DMEM) (Gibco; Thermo Fisher Scientific, Inc., Waltham, MA, USA), supplemented with 10% fetal bovine serum (Zhejiang Meisen Cell Technology Co., Ltd., Hangzhou, China) and 1% penicillin-streptomycin solution (Seven Innovation Biotechnology Co., Ltd., Beijing, China) at culture incubator (37 °C, 5% CO2, and 95% humidity).

### 2.2. Cell Viability Assay

The cell viability was tested by a Cell Counting Kit-8 (CCK-8) purchased from Biosharp (Hefei, China). C2C12 cells were plated in a 96-well plate at 2000 cells per well. To prevent evaporation, 100 μL of PBS solution was added to the outermost wells of the 96-well plate, with six replicate wells for each treatment concentration. After 24 hours of drug treatment, the supernatant was removed, and the cells were washed 1~2 times with PBS. A 10% CCK-8 solution was prepared and added to the wells, followed by incubation at 37 °C for 2 hours. After incubation, a multifunctional microplate reader was used to determine the absorbance of the samples at 450 nm, and the measurement results were recorded and exported.

### 2.3. ROS Assay

Cells were seeded into a 24-well plate and incubated with the drug after 24 hours of culture. The cells were washed twice with PBS (at room temperature), and then 500 μL of PBS containing 10 μM H2DCFDA (Biosharp, Hefei, China) was added to each group of cells, followed by incubation in a cell culture incubator at 37 °C for 30 minutes in the dark. After incubation, the cells were washed 1-2 times with PBS to fully remove any H2DCFDA that had not entered the cells. The cells were then collected, resuspended, and mixed evenly in PBS. The solution was evenly dispensed back into the microplate, with 100 μL per well, and the remaining cell suspension was used for counting. The microplate was placed in a microplate reader to determine the fluorescence value at an excitation wavelength of 488 nm and an emission wavelength of 525 nm.

### 2.4 Superoxide Dismutase Activity Assay

The protein concentration of the samples was normalized using the BCA method, followed by determination using Superoxide Dismutase (SOD)Activity Assay Kit (WST-1 Method) (Beijing Solaibao Technology Co., Ltd., Beijing, China). According to the instructions provided by the kit, reagent 2 and reagent 4 working solutions were prepared and added to the samples and other solutions in sequence. After mixing thoroughly, the mixture was incubated in a water bath at 37 °C for 30 minutes. The 96-well plate was then placed in Spark Microplate Reader (Tecan (Shanghai) Trading Co., Ltd., Shanghai, China) to measure the absorbance of the samples at 450 nm.

### 2.5 Glutathione Peroxidase Activity Assay

Before measurement, reagents from the GSH-Px assay kit (Geruisi Biotechnology Co., Ltd., Suzhou, China) were incubated in a water bath at 25 °C for 30 minutes. According to the instructions, reagents were added to the assay tube and control tube. After the reaction was completed, the solution was evenly dispensed into the enzyme-linked immunosorbent assay (ELISA) plate, and the absorbance of the samples was measured at 412 nm.

### 2.6 MDA Assay

To a centrifuge tube, 300 μL of working solution and 200 μL of the sample were added, followed by incubation in a water bath at 90~95 °C for 30 minutes. After incubation, the tube was placed on ice to cool to room temperature, then centrifuged at 12000 rpm for 10 minutes at 25 °C. Subsequently, 200 μL of the supernatant was transferred to a 96-well plate, and the absorbance of the sample was measured at 532 nm and 600 nm, respectively.

### 2.7 RNA-seq and Analysis

Total RNA was extracted using TRIzol Reagent (Invitrogen, Thermo Fisher Scientific, Inc., Waltham, MA, USA). RNA purification, genomic DNA removal, RNA quality inspection, and sequencing were all handled by Wuhan Yingzi Gene Technology Co., Ltd.

During the data analysis, FastQC (v0.11.8) [17] was utilized for data quality control. To ensure the reliability of data analysis, raw data needed to be filtered. Trimmomatic [18] was used to filter out low-quality reads from the raw data. The Q20, Q30, and GC content of the clean reads were calculated. All subsequent analyses were based on these clean reads. lncRNA/mRNA data were aligned to the mouse reference genome using hisat2[19]. Samtools [20] was used to collect alignment information, and StringTie [21] was applied to assemble transcripts for each sample individually.

### 2.8 Dual-luciferase assay

After plasmid extraction using an Endo-Free Plasmid Mini Kit I (Guangzhou Feiyang Biological Engineering Co., Ltd., Guangzhou, China), the wild-type, mutant, empty vector, mimics, and mimics NC were co-transfected using Lipofectamine^®^ LTX & PLUTM Reagent (Invitrogen, Thermo Fisher Scientific, Inc., Waltham, MA, USA) for 48 hours. The Dual-Luciferase^®^ Reporter Assay System (Promega (Beijing) Biotechnology Co., Ltd., Beijing, China) was then used to assay the activities of firefly luciferase and renilla luciferase.

### 2.9 Statistical Analysis

The data presented in this study were obtained from three independent biological replicate experiments. Statistical analysis and visualization were performed using GraphPad Prism. Significance was determined using one-way ANOVA followed by a t-test, with *p* < 0.05 considered statistically significant.

## 3. Results

### 3.1. Effect of PA and BS on C2C12 Cells

The analysis of the effect of different concentrations of PA on the viability of C2C12 cells is presented in Figure 1.

**Figure 1.**
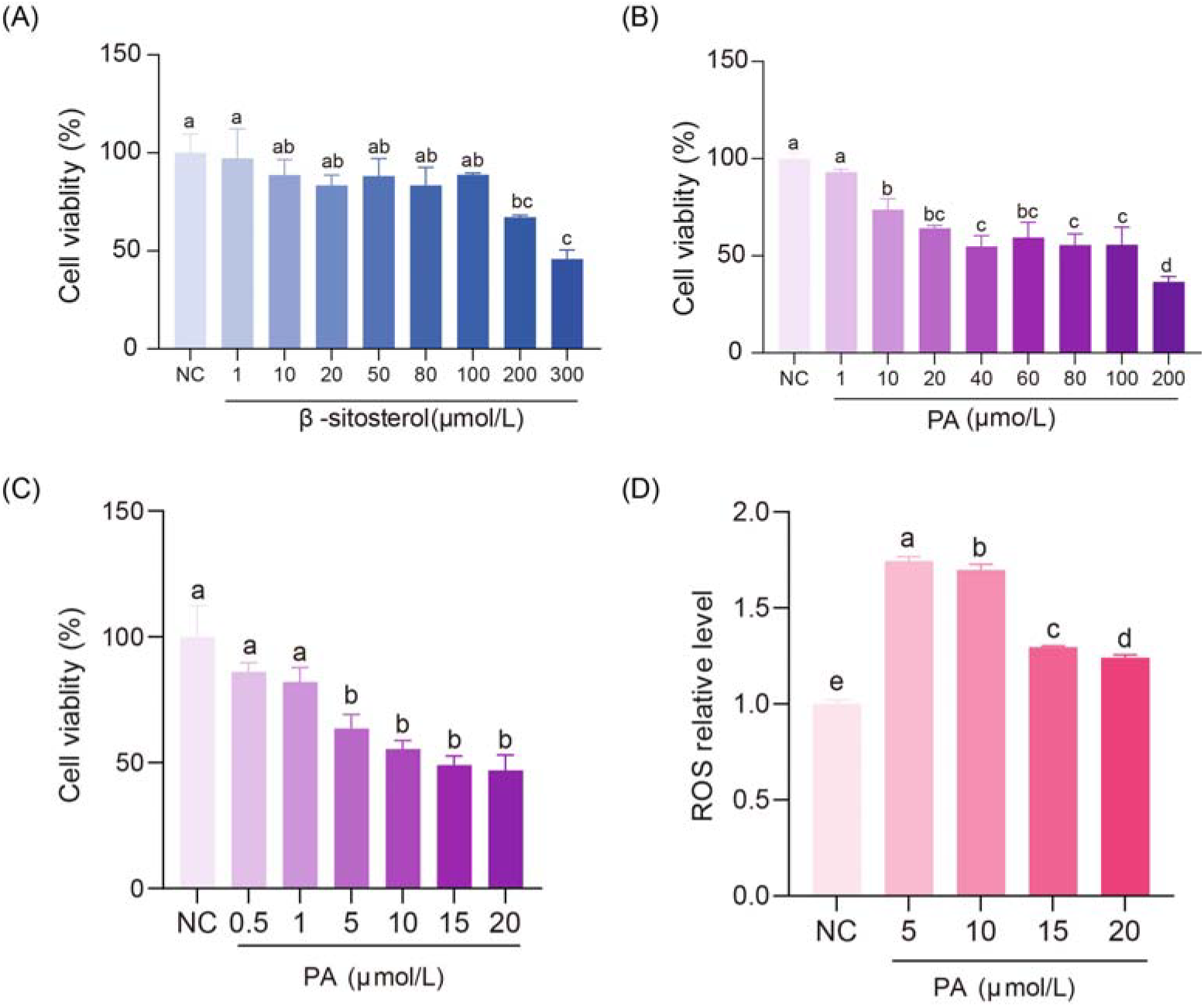
Effect of PA and β-sitosterol on cell viability and ROS relative level. (A) The C2C12 cells were treated with 1~300 μmol/L β-sitosterol for 24 h, and then the cell vitality was assayed using a CCK-8 assay. (B) The C2C12 cells were treated with 1~200 μmol/L PA for 24 h, and then the cell vitality was assayed using a CCK-8 assay. (C) The C2C12 cells were treated with 0.5-20 μmol/L PA for 24 h, and then the cell vitality was assayed using a CCK-8 assay. (D) Effect of PA on ROS generation in C2C12 cells. Different letters represent a statistically significant difference (*p* < 0.05).

Figure.1A shows that when BS concentration is less than or equal to100 μmol/L, compared with the control group (NC group) with 100% cell viability, there is no significant effect on cell viability (*p* < 0.05). Therefore, a concentration of 100 μmol/L was chosen for subsequent experiments in this study.

As shown in Figure 1B, compared to the control group (NC group) with 100% cell viability, PA concentrations of 10 μmol/L or higher have a significant decrease the cell viability (*p* < 0.05). To further explore the optimal modeling concentration, additional investigations were conducted within the selected concentration range. Figure 1C illustrates the effect of PA concentrations ranging from 0.5 to 20 μmol/L on the viability of C2C12 cells. Compared to the control group (NC group) with a cell viability of 100%, PA concentrations of 5 μmol/L or higher significantly decrease the cell viability (*p* < 0.05), with no significant difference observed between 5 and 20 μmol/L.

Oxidative stress is characterized by the excessive accumulation of ROS. Figure 1D illustrates the effect of different concentrations of PA on ROS levels. The ROS experiment results indicate that, compared to the control group (NC group), stimulation with 5~20 μmol/L of PA can induce significant ROS production in C2C12 cells (*p* < 0.05), suggesting that PA stimulation is capable of causing oxidative stress in cells.

Based on the combined results of cell viability and ROS production, PA concentrations of 5 μmol/L was ultimately selected for modeling oxidative stress in C2C12 cells.

### 3.2. BS repair. PA-Induced Oxidative Damage in C2C12 Myoblasts

Figure 2 demonstrates that following 24 hours of oxidative stress modeling in C2C12 cells induced by 5 μmol/L of PA, BS significantly restored five related indices (CCK8, ROS, SOD, GPx, and MDA). This suggests that under conditions of mild oxidative stress, the self-repair capability of cells is not completely destroyed, and BS can effectively support the reconstruction of the cellular antioxidant system and damage repair.

**Figure 2.**
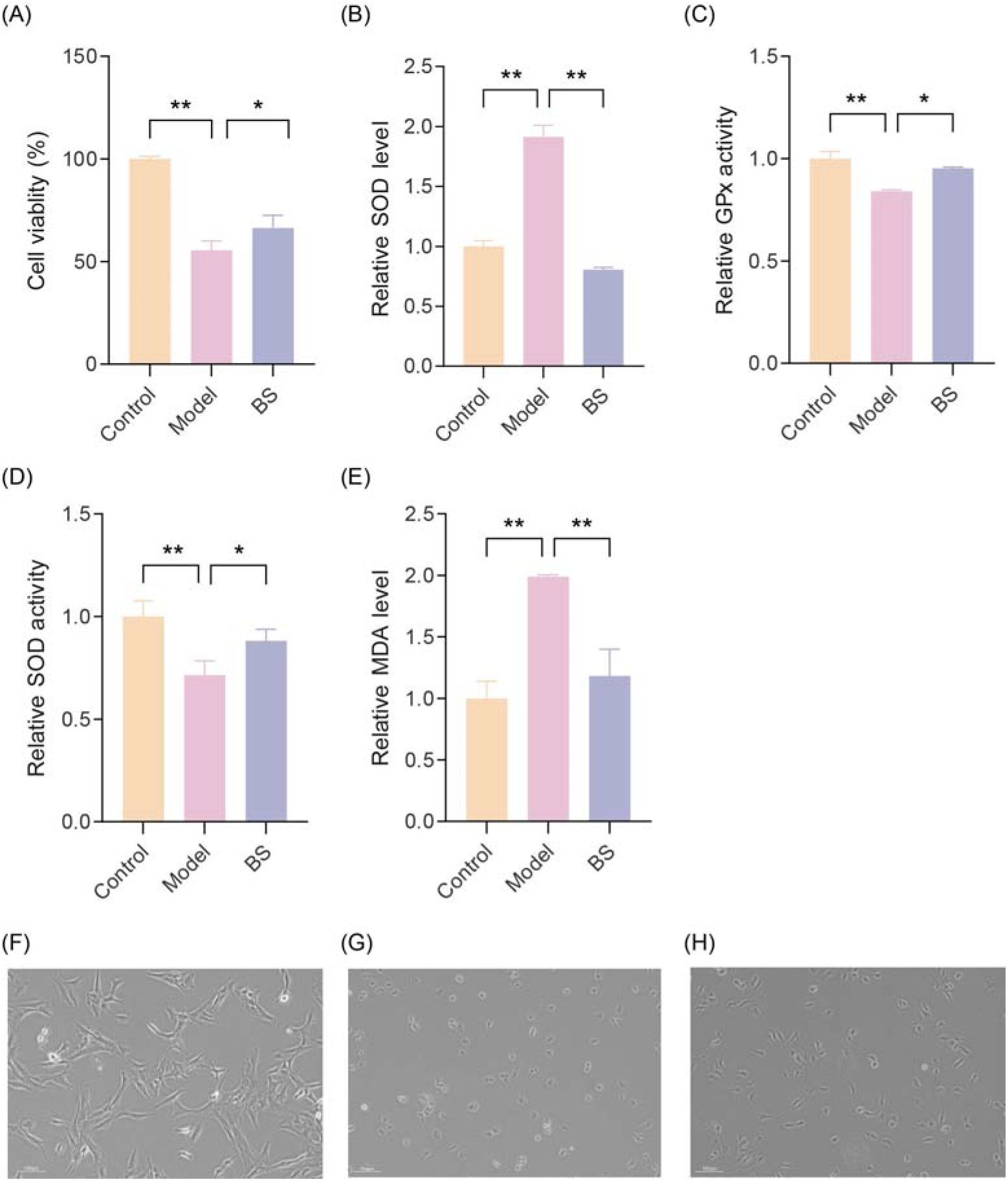
Changes of related indexes after BS treatment with low concentration PA molding. (A) The effect of BS on the cell viability of C2C12 cells after oxidative damage. (B) The effect of BS on the ROS relative level of C2C12 cells after oxidative damage. (C) The effect of BS on the GPx activity of C2C12 cells after oxidative damage. (D) The effect of BS on the SOD activity of C2C12 cells after oxidative damage. (E) The effect of BS on the MDA relative level of C2C12 cells after oxida-tive damage. (F) Control group was cultured normally, and no drugs were given. (G) PA modeling group. (H) Rescue with BS after PA modeling. “*” represents *p* < 0.05, and “**” represents *p* < 0.01. Images were captured under a 10× microscope, with a scale bar representing 100 μm.

Under normal culture conditions, that is, without adding any drugs, C2C12 cells grow adherent to the substrate, exhibiting a spindle-shaped morphology with radiating branches and long fibers extending in multiple directions. (Figure 2F). Upon PA treatment, the cell morphology in the PA group gradually transitions from spindle-shaped to rounded compared to the control group (Figure 2G). However, upon subsequent recovery treatment with BS, the cells gradually regain their spindle shape (Figure 2H).

### 3.3. RNA-seq Analysis

Based on RNA-seq data, the gene expression profiling of control groups and PA groups, PA groups and BS groups were further analyzed. The RNA sequencing results of each group were analyzed, and the genes with |log2foldchange| > 1 and *p* < 0.05 were selected for statistical analysis. As shown in the volcanic map (Figure 3A, B), the expression of mmu-miR-1291 increased under the condition of oxidative stress, and after BS treatment, the expression of mmu-miR-1291 increased.

**Figure 3.**
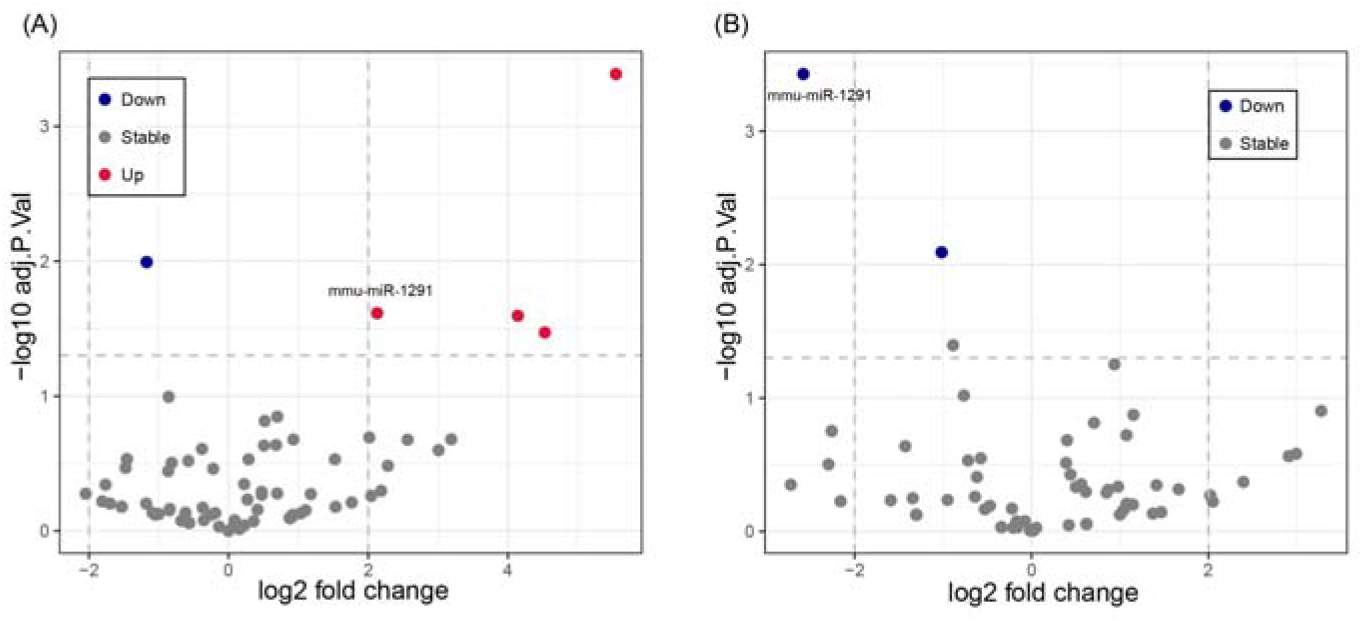
(A) The volcano plot analysis of differentially expressed miRNAs in control and PA groups of C2C12 cells by RNA-seq. (B) The volcano figure analysis of differentially expressed miRNAs in PA and BS groups of C2C12 cells by RNA-seq.

### 3.4. Prediction and Enrichment Analysis of Target Genes

The target gene bound by mmu-miR-1291 was predicted by using the online open database data of TargetScan, miRWalk and miRDB. According to the analysis results of the three databases, 309 candidate target genes were screened (Figure 4.). The predicted target genes were subjected to enrichment analysis and *Wsb1* was enriched to the entries shown in Figure 5. And *Wsb1* was found to be predicted in all three databases and ranked high in both TargetScan and miRDB (Table1, 2).

**Table 1.**
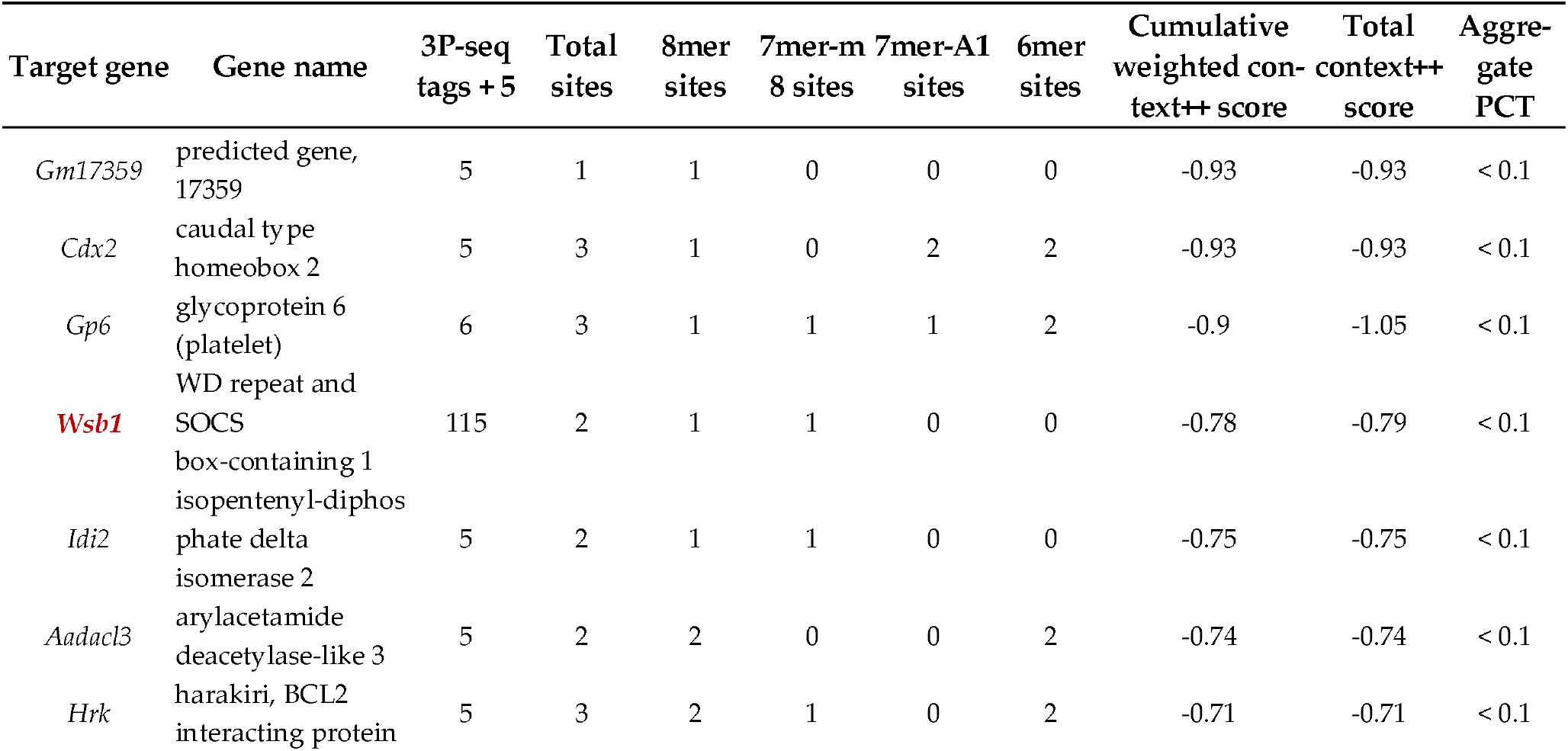
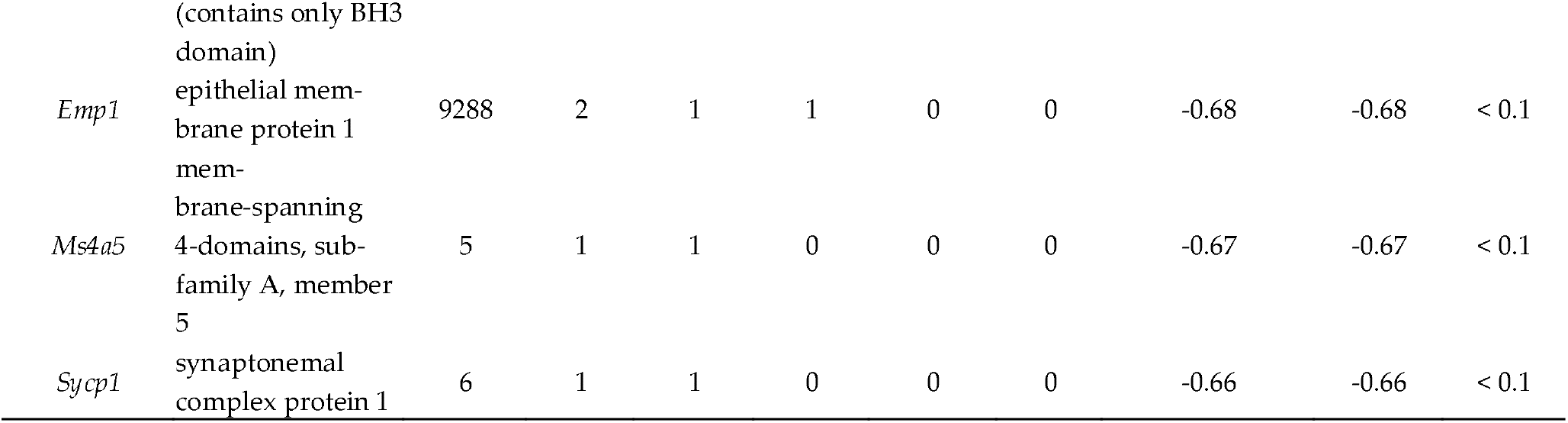
TargetScan predicts specific information of target genes.

**Table 2.** miRDB predicts specific information of target genes.

| Target rank | Target score | Gene symbol | Gene description |
| --- | --- | --- | --- |
| 1 | 98 | *E2f4* | E2F transcription factor 4 |
| 2 | 97 | *Dnajc16* | DnaJ heat shock protein family (Hsp40) member C16 |
| 4 | 96 | *Itpr1* | inositol 1,4,5-trisphosphate receptor 1 |
| 5 | 96 | ***Wsb1*** | WD repeat and SOCS box-containing 1 |
| 6 | 94 | *Ctsc* | cathepsin C |
| 7 | 94 | *Phf20* | PHD finger protein 20 |
| 8 | 93 | *Lrrtm2* | leucine rich repeat transmembrane neuronal 2 |
| 9 | 93 | *Birc6* | baculoviral IAP repeat-containing 6 |
| 10 | 93 | *Sycp1* | synaptonemal complex protein 1 |
| 11 | 92 | *Adam12* | a disintegrin and metallopeptidase domain 12 (meltrin alpha) |

**Figure 4.**
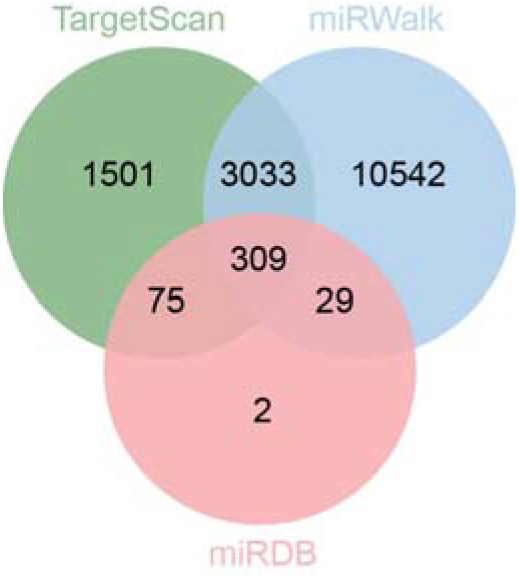
Venny diagram showing TargetScan, miRWalk and miRDB prediction of mmu-miR-1291 target genes.

**Figure 5.**
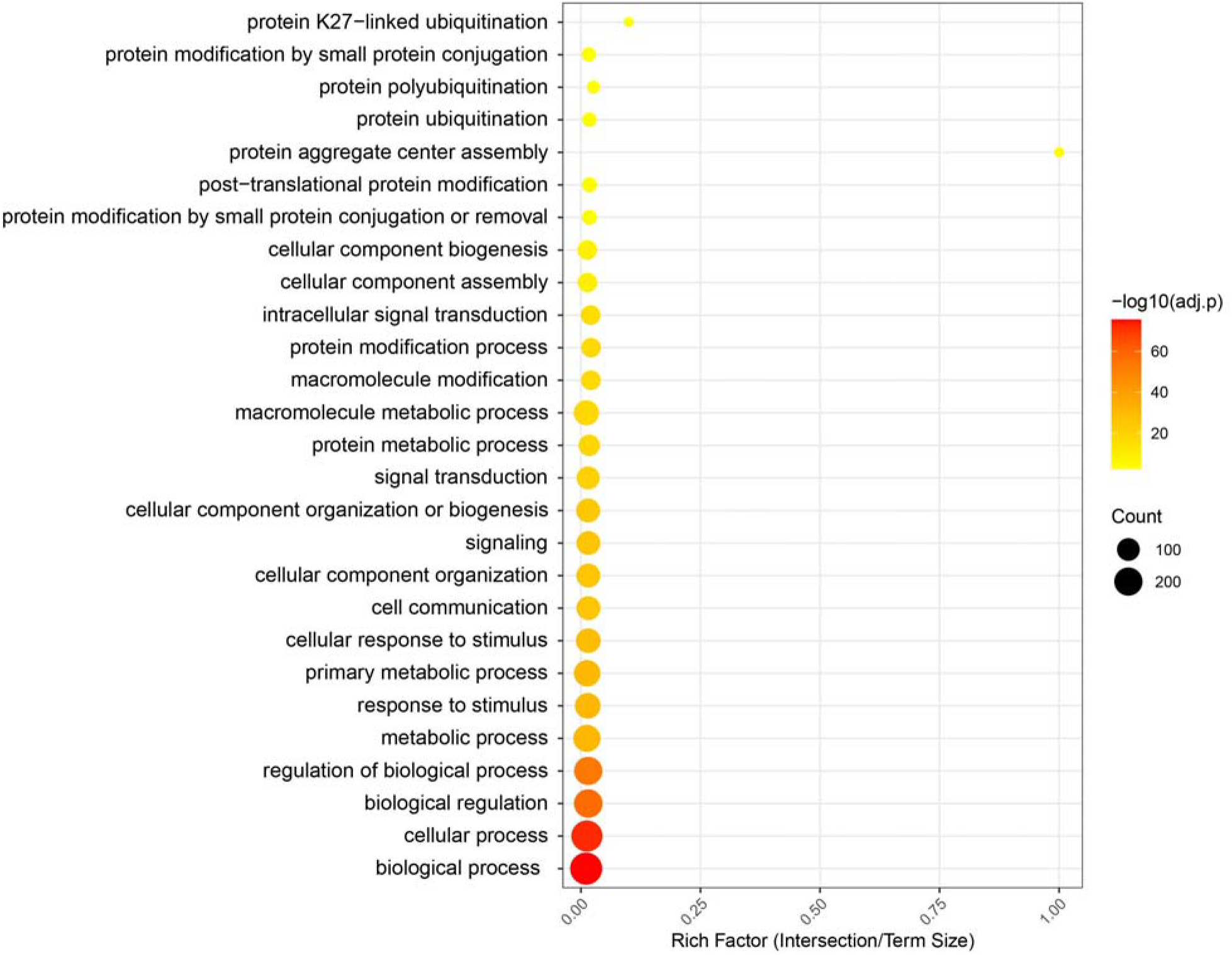
Gene ontology enrichment analysis

### 3.5. Assay of Wsb1 3’-UTR Activity Using the Dual-Luciferase Reporter Gene System

After co-transfection of C2C12 cells with *Wsb1* wild-type, mutant, empty vector, and mmu-miR-1291 mimics or mimics NC for 48 hours, the activities of firefly luciferase and renilla luciferase were measured using a fluorometer. The results are presented in Figure 15. The data revealed a significant decrease in fluorescence in the co-transfected group with mmu-miR-1291 mimics and *Wsb1* 3’-UTR WT compared to the co-transfected group with mimics NC and *Wsb1* WT, showing a significant difference (*p* < 0.01). However, there was no significant change in fluorescence between the co-transfected group with mmu-miR-1291 mimics and *Wsb1* 3’-UTR WT compared to the co-transfected group with mimics NC and *Wsb1* MUT.

**Figure 15.**
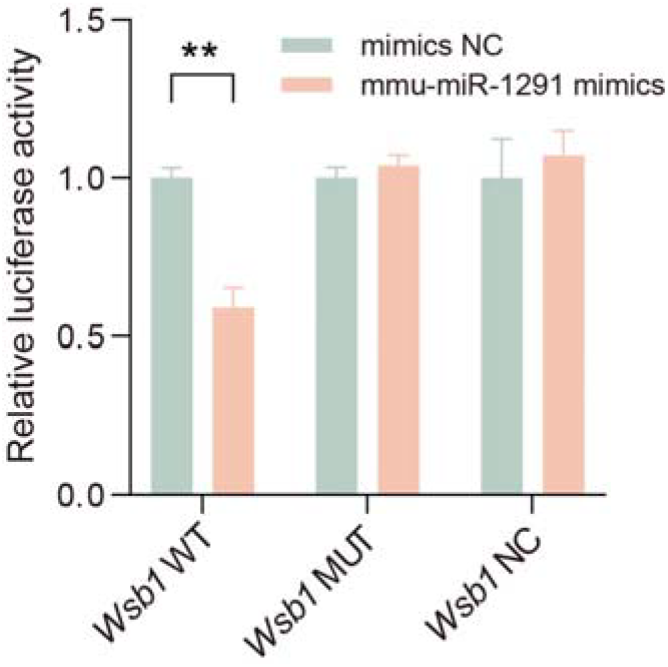
Assay of Dual-Luciferase Activity.

## 4. Discussion

Oxidative stress is caused by an imbalance between the production of ROS and the antioxidant defense system of the body. Excessive ROS can damage cellular macromolecules, such as lipids, proteins, and nucleic acids, triggering oxidative stress and resulting in cell injury. Skeletal muscle is the largest tissue in mammals by mass, crucial for maintaining overall health and physiological functions. It is the main source of animal protein, and its growth and development have a direct impact on meat production and quality traits. Skeletal muscle is rich in mitochondria, which are the primary sites for ROS generation. Therefore, maintaining the health of skeletal muscle cells is crucial for promoting animal growth and development and improving production performance.

Palmitic acid is the most prevalent saturated free fatty acid. When overloaded in non-adipose cells, it can lead to lipotoxicity and cell apoptosis [22]. Palmitic acid-induced oxidative stress models are widely used to investigate cellular responses to oxidative stress. BS, as a key active component of plant sterols, is one of the richest plant sterols and is a safe, natural, green, and effective animal nutritional supplement with multiple pharmacological activities, including anti-inflammatory, antioxidant, cholesterol-lowering, blood sugar-lowering, and immune-regulating effects [23]. BS is abundantly present in oilseed plants, medicinal plants, grains, fruits, and other sources [24]. BS, as a bioactive substance that can be extracted from agricultural by-products, has a reparative effect on oxidative damage in C2C12 mouse myoblasts.

This study uses PA to stimulate C2C12 cells, inducing excessive ROS production to explore whether BS can improve the oxidative damage in the cells. Oxidative damage inside and outside the cell under oxidative stress is often assessed through changes in a series of markers. The increase in MDA content, changes in SOD and GPx activity, and the rise in ROS levels are widely regarded as important biological markers of cellular oxidative damage. Among these, MDA is the final product of lipid peroxidation, and its increased level reflects the degree of oxidative damage to the cell membrane lipids. SOD and GPx, as major antioxidant enzymes, show changes in activity that indicate the anti-oxidant capacity of the cells. Oxidative stress is also accompanied by a significant increase in ROS content. The accumulation of ROS further induces damage to proteins, DNA, and lipids, affecting cell function and survival.

The experimental results show that PA stimulates C2C12 cells, causing excessive ROS production and significantly inhibiting the activity of antioxidant enzymes within the cells, promoting cell apoptosis. The degree of damage to C2C12 cells induced by oxidative stress at different concentrations of PA varies significantly, which in turn affects the recovery effect of the drug. Under the 5 μM PA, the oxidative stress is relatively mild, and the antioxidant system of the cells has not been completely destroyed, allowing significant recovery within 24 hours of drug treatment. However, under the 10 μM PA the oxidative stress is more severe, and the significant increase in ROS levels and suppression of cell vitality require a longer period for repair. Although the drug treatment for 24 hours significantly restores the activity of antioxidant enzymes (SOD, GPx) and reduces lipid peroxidation products (MDA), the complete recovery of ROS clearance and cell viability (CCK8) is only achieved after 48 hours and 72 hours, respectively. Oxidative stress induced by 10 μM PA is more severe, and the extensive accumulation of ROS may damage mitochondrial function, inhibit cell proliferation, and induce a series of consequences such as lipid peroxidation and protein oxidation. Therefore, the recovery of CCK8 and ROS exhibits a delayed time response. This indicates a time-dependent relationship between oxidative stress intensity and cellular recovery, further highlighting the cumulative effect of drug action and the dynamic nature of the repair process.

During the sequencing data analysis, we identified differentially expressed mmu-miR-1291. So far, there have been no reports have been published domestically or internationally regarding the role of mmu-miR-1291 in inflammation or oxidative stress. Current studies on mmu-miR-1291 have primarily focused on cancer cells, including pancreatic cancer [42], prostate cancer [43], and esophageal squamous cell carcinoma [44], suggesting that it may serve as a novel therapeutic target in oxidative stress. To further investigate mmu-miR-1291, we conducted target gene prediction using three online databases and identified 309 potential target genes.

After screening these candidates, we selected *Wsb1* for experimental validation. *Wsb1* is a substrate recognition component of the Cullin-RING E3 ubiquitin ligase, which targets chromatin-bound methylated RelA (a subunit of NF-κB) for polyubiquitination and proteasomal degradation, thereby terminating NF-κB transcriptional activity and facilitating inflammation resolution [45].

These enriched GO entries correlate with the functions of Wsb1 in the biological processes of cells. *Wsb1* acts as a key player in cellular homeostasis by regulating cellular response to different stimuli, such as oxidative stress. Oxidative stress causes damage to intracellular structure and function through excess free radicals generated by ROS, and *Wsb1* may help cells to remove damaged proteins through its ubiquitination function, regulating intracellular homeostasis and mitigating the damage caused by oxidative stress.

To verify the regulatory interaction between mmu-miR-1291 and *Wsb1*, we employed a dual-luciferase reporter assay. We co-transfected psiCHECK-2 recombinant plasmids containing either the wild-type or mutant *Wsb1* 3’-UTR binding sites with miRNA mimics (mimics/mimics NC) into C2C12 cells. Luciferase activity assays revealed that mmu-miR-1291 specifically recognized the *Wsb1* 3’-UTR region, leading to a significant downregulation of luciferase reporter activity (*p* < 0.01). These results indicate that mmu-miR-1291 mediates post-transcriptional regulation by binding to the *Wsb1* 3’-UTR target site. This study is the first to experimentally validate *Wsb1* as a potential target gene of mmu-miR-1291.

## 5. Conclusions

BS exhibited a significant restorative effect on PA-induced oxidative damage in C2C12 cells. And a set of miRNAs that were significantly differentially expressed under oxidative stress conditions was identified. Through target gene prediction, a key miRNA potentially involved in oxidative stress regulation was selected. Additionally, differential expression and enrichment analyses led to the identification of a series of mRNAs affected by oxidative stress. Based on miRNA target prediction, the *Wsb1* gene was selected for further investigation. A dual-luciferase reporter assay confirmed that this miRNA directly regulates *Wsb1*. This suggests that the identified miRNA may influence the oxidative stress process by regulating *Wsb1* expression and potentially be involved in modulating the NF-κB signaling pathway.

## Author Contributions

Conceptualization, S.J., D.B. and Y.B.; writing—original draft preparation, S.J. and D.B.; writing—review and editing, S.J., Y.W. and D.B.; data curation, Y.W. and S.J.; supervision, Y.B. All authors have read and agreed to the published version of the manuscript.

## Data Availability Statement

The data presented in this study are available on request from the corresponding author.

